# Discordance in Genotypic and Phenotypic anti-tuberculosis drug susceptibility results: time to reconsider critical concentration

**DOI:** 10.1101/2024.07.08.602571

**Authors:** Arti Shrivas, Sarman Singh, Jitendra Singh, Prem Shankar, Payal Soni, Syed Beenish Rufai, Anand Maurya, Shashank Purwar

## Abstract

**Objective:** To correlate *rpoB* mutations found on the sanger sequencing in *Mycobacterium tuberculosis* (MTB) isolates with Minimum Inhibitory Concentrations (MICs) to the rifampicin.

**Methods:** We assessed the minimum inhibitory concentrations (MICs) for 151 archived clinical MTB isolates that were determined phenotypically susceptible to RIF (101;66.89%) and remaining fifty (50;33.11%) were resistant to RIF by BACTEC MGIT SIRE DST. MIC values were determined using colorimetric redox indicator (Resazurin/REMA) method and results were correlated with *rpoB* gene mutations associate with rifampicin resistance found.

**Results:** Comparing the MIC and critical concentration, we found that 15 of these 101 (14.85%) isolates were misclassified by MGIT-960 as sensitive at standard critical concentration (1.0µg/mL) though these were found to have low-level RIF resistance by CRI assay (MIC 0.50µg/mL to 1.0µg/mL) and sanger sequencing. We found that all of 15 isolates contained non-synonymous mutations, the commonest being the *Ile572Phe* (7, 46.66%), followed by *Leu533Pro* (3, 20.0%), *His526Leu* (2, 13.33%), *His526Asn*+*Ile572Phe* (1), *Asp516Tyr* (1), and *Leu533Pro*+*Pro564Arg* (1). These mutations are reported to confer low-level RIF resistance. But we did not find any mutation at MIC ≤0.25μg/mL.

**Conclusion:** We found that a significant number of MTB isolates have phenotypic and genotypic discordance. Taking 1.0µg/mL of rifampicin as critical concentration, isolates from approximately 15% patients are misidentified as susceptible to rifampicin, even when these strains carry low level drug resistance conferring mutations and have potential to develop clinical MDR-TB.

## Introduction

*Mycobacterium tuberculosis* (MTB) is the causative agent of tuberculosis (TB). Drug resistance TB has become a major hurdle in the treatment and control of TB around the globe, threatens the progress made so far by the TB elimination programs (1). Patients infected with strains resistant to isoniazid (INH) and rifampicin (RIF) the two most effective first-line anti-TB drugs called multidrug-resistant (MDR) TB, are practically refractory to standard first-line treatment (2). The past decade has seen the increment of MDR-TB around the globe. The number of primary MDR-TB remained relatively decreased from 3.90% in 2015 to 3.10% in 2020 but started increasing in 2021 and reached to 3.60%, despite COVID-19 epidemic which caused under-reporting (3). India notified 24.20 lakh MDR cases in the year 2022; an increase of 13.0% as compared to 2021 and reported a total number of 63,801 MDR cases (4).

Rifampicin (RIF) is an important constituent of first- and second-line treatment. This drug acts against actively growing and slowly metabolizing (non-growing) bacilli (5). However, prior to start of MDR-TB treatment it is necessary to carry out rapid drug susceptibility testing to avoid the misuse of ineffective drugs. In accordance with WHO guidelines, detection of MDR-TB requires testing for drug resistance using rapid molecular tests, culture methods or sequencing technologies (2,6,7). However, conventional culture based drug susceptibility (DST) also known as phenotypic methods are considered to be the ‘gold standard’ for determining MTB drug resistance (8). Various molecular assays like GeneXpert MTB/RIF and Line Probe Assay/MTBDR*plus* are being progressively applied to detect rifampicin resistance (RR) or MDR.

Resistance to RIF is mainly associated with the mutations in the *rpoB* gene of MTB. Furthermore, more than 95% of MTB clinical isolates that are resistant to RIF have a mutation in an 81bp ‘hot spot’ region of *rpoB* known as the rifampicin resistance–determining region (RRDR) between codons 507–533 (9). Position of these mutations within the *rpoB* gene are associated with different levels of phenotypic resistance to rifampicin. Some mutations are associated with a high-level of resistance to RIF while others are known to confer only low level resistance (10).

Although, the molecular tests have been considered to be highly accurate initially but now some of the clinically relevant mutations conferring resistance are being reported to be missed by these assays because of location of these mutations outside the targeted region (11,12).

Similarly, the culture-based methods are also reported to miss clinically relevant RIF resistance in some isolates despite having genetic mutations in *rpoB* region of MTB. The MTB isolates with undisputed mutations in hot-spot region (HSR) of *rpoB* gene are readily detected by phenotypic DST methods (13). However, there are frequent discordant results between phenotypic and genotypic assays as shown in various studies (14–17).

Low-level resistance to RIF is clinically significant as patients infected with *M. tuberculosis* strains with disputed *rpoB* mutations often fail treatment or relapse (18,19). Several recent studies target to identify characteristics of the low-level RR-TB patients worldwide to understand the resistance pattern found, search for similar strains and identify possible implications for treatment recommendations (20–23).

It has been previously reported in many countries that the mutations associated with low level resistance which are not detected by any of the two WHO endorsed methods i.e. phenotypic and molecular DST. However, there is scarcity of data from India regarding the presence of these mutations.

Limited research has raised concerning the assay performance in patient care settings with respect to specificity, sensitivity, indeterminate results, inefficient detection of rifampicin mono-resistance due to mutations present inside or outside RRDR region and certain disputed mutations. It is felt that by the continued use of WHO notified critical concentration of RIF, we might be missing several cases of potential RIF resistance, and these patients are administered with the ineffective anti-tuberculosis drug regimen. Therefore, in present study we aimed to determine the minimum inhibitory concentration (MIC) in RIF sensitive and RIF resistant isolates following the standard protocols, analysed the genetic mutations in *rpoB* by gene target sequencing and compared the MIC with various genetic mutations.

## Material method

### Study setting

The study was carried out from April 2020 to September 2021. The study was approved by the Institutional human Ethics Committee of the All India Institute of Medical Sciences Bhopal (IHECPGRPHD063).

### Clinical isolates

A well characterized and anonymised 151 MTB culture isolates were randomly selected from the repository maintained in the accredited TB research laboratory, at the All India institute of Medical Sciences, New Delhi, India. The drug susceptibility pattern of these isolates was previously identified by BACTEC MGIT960 SIRE DST system. After characterization, these isolates were stored in minus 80°C with *in-house* culture cataloguing until further use. These MTB isolates were freshly sub-cultured for this work on Lowenstein Jensen (LJ) medium and in MGIT 960 culture tubes before being used for further phenotypic and genotypic characterization.

### Phenotypic DST

Drug susceptibility testing (DST) was performed for all the above 151 MTB isolates by using BACTEC MGIT 960 SIRE Kit method (Becton-Dickinson, USA). A stock solution was prepared by reconstituting the lyophilized drugs by adding 4.0mL of sterile distilled water to each vial to make a stock solution of Streptomycin (83.0µg/mL), (Isoniazid) 8.30µg/mL, (Rifampicin) 83.0µg/mL and (Ethambutol) 415.0µg/mL. All four drugs were used at a final concentration of 1.0µg/mL (STR), 0.1µg/mL (INH), 1.0µg/mL (RIF) and 5.0µg/mL (EMB) in MGIT tube respectively Once reconstituted the drugs were used for the test immediately. A susceptible growth control isolate, H37Rv (American Type Culture Collection, 27294). The remaining antibiotics solutions were stored at -20°C for future use. Preparation of the inoculum, as well as inoculation and incubation were performed as per manufacturer instructions (Becton Dickinson BACTEC MGIT 960 system, BD, USA)(24)

### Determination of minimum inhibitory concentration for rifampicin

Minimum inhibitory concentration (MIC) of Rifampicin (RIF) was determined of all the 151 well characterized *M. tuberculosis* laboratory isolates by colorimetric redox indicator assay using resazurin salt as redox indicator. Briefly, 100 μL of 7H9-S broth (Middlebrook 7H9 supplemented with 0.1% Casitone, 0.5% glycerol, and 10% OADC [oleic acid, albumin, dextrose, and catalase]; Becton-Dickinson) was dispensed in each well of a sterile flat-bottom 96-well plate. Working solutions of the drug was initially prepared at four times the final higher concentration in 7H9-S broth. One hundred microliters of the working drug concentration (rifampicin 8.0 μL/mL) were added to the wells containing 7H9-S broth and the drug was then further serially diluted two-fold to a final concentration of rifampicin (0.0625 μL/mL)(25).

An inoculum turbidity of McFarland standard number 1.0 was prepared from 7H9-S broth, diluted (1:20) in 7H9-S broth for test. One hundred microliters of inoculum was added to the drug-free and drug-containing wells. A sterile control and a growth control for each isolate were also included. To prevent evaporation during incubation, sterile water was added to all perimeter wells. The plates were incubated at 37°C after being sealed in a plastic bag. A working solution of resazurin salt (30 μL of 0.02% concentration) was inoculated after 7 days of incubation into each micro well. After overnight incubation, plates were then read the next day, a total of 9 days turnaround time for results interpretation. A colour change from blue to pink denoted a positive reaction (reduction of resazurin to resorufin) confirming drug resistance and growing *M. tuberculosis* cells. The MIC was defined as the lowest concentration of drug that prevented colour change.

The isolates which showed MIC value either equal to or higher than 0.50μg/mL (Breakpoint) were considered as resistant or uninhibited. For test interpretation, the colour was compared to the colour present in the growth control well. The first well with complete inhibition of growth was considered as MIC for that isolate. Confirmation of cells viability was done by inoculation of those cells on agar plates. The range of drug concentration we used was 2.0µg/mL to 0.0625µg/mL. Each isolate was tested in duplicates in the same plate.

### DNA Extraction

Genomic DNA of twenty-four *M. tuberculosis* isolates was extracted from MTB cultures grown on L-J medium using chloroform iso-amyl alcohol (CI) method as mentioned previously (Reference) and DNA yields were measured by Qubit fluorometry to confirm purity (26).

### Multiplex-PCR assay

Prior to sequencing, a well-established multiplex polymerase chain reaction (mPCR) assay was utilized for the identification of *Mycobacterium tuberculosis* Complex (MTBC). The extracted DNA were identified with genus and species-specific primers. Primers were selected to amplify the regions of heat shock protein-65 (hsp-65), early secretory antigenic target-6 (esat-6) and internally transcribed sequence (*its*) to detect Mycobacterium genus, M. tuberculosis complex (MTBC) species and M. avium complex (MAC) species respectively (27).

PCR products were electrophoresed on 1.8% agarose gel and visualized under UV trans illuminator results were interpreted by *M. tuberculosis* specific bands at 441bp (*hsp-65*) and 320bp (*esat-6*), and 126bp (*its*).

### Primer Designing and sequencing

The PCR primer sets targeting inside and outside RRDR region of *rpoB* gene, *M. tuberculosis* H37Rv (GenBank accession no: NC_000962) were designed by Primer3software (http://primer3.ut.ee/) and custom synthesized from Eurofins Genomics India Pvt. Ltd. Bangalore, India. The sequencing primers used were forward: 5’-ATG ACC ACC CAG GAC GTG-3’ and reverse: 5’-ACA CGA TCT CGT CGC TAA CC-3’. A PCR assay with sequencing primers was also performed as a quality control check on DNA extracts to assess sequencing of *rpoB* gene and quantitation of *Mycobacterium tuberculosis* DNA.

A total of twenty-four representative isolates (RIF Sensitive 20; RIF Resistant 04) were sequence confirmed by Sanger sequencing. These isolates were selected based on the MIC results to determine the level of phenotypic resistance with mutations associated with *rpoB* gene in *Mycobacterium tuberculosis.* PCR products were purified by using QIAquick PCR purification kit protocol. The purified amplicons were commercially sequenced from Anuvanshiki (OPC) Pvt Ltd., New Delhi, using automated sequencer (Applied Biosystem).

### Sequencing result analysis

The obtained sequences were processed and analysed through MEGA X for mutational study. Mutations in *rpoB* gene sequences were compared with wild type sequences of H37Rv strain of *M. tuberculosis* using ClustalW tool (http://www.ebi.ac.uk/Tools/msa/clustalw2) and again performed multiple sequence alignments (MSA) using ClustalW tool.

## Results

A total of 151 well characterized MTB culture isolates were randomly selected for this study. Among these, 50 (33.11%) were resistant to rifampicin (RIF), singly or in combination with other drugs, and 101(66.89%) isolates were phenotypically sensitive to RIF. The drug susceptibility pattern of these isolates is given in table (1).

**Table 1.** Phenotypic drug susceptibility patterns of the MTB culture isolates (N=151)

| S. No. | RIF susceptibility status | Sensitive/Resistance to drugs | No. (%) |
| --- | --- | --- | --- |
| 1. | RIF sensitive isolates<br>(n=101; 66.88%) | STR+INH+RIF+EMB Sensitive | 70 (69.3%) |
|  |  | INH Mono-Resistant | 20 (19.8%) |
|  |  | STR Mono-Resistant | 3 (3%) |
|  |  | EMB Mono-Resistant | 2 (2%) |
|  |  | INH+EMB Resistance | 5 (4.9%) |
|  |  | INH+STR Resistant | 1 (1%) |
| 2. | RIF resistant isolates<br>(n=50; 33.11%) | RIF+INH Resistant | 20 (40%) |
|  |  | RIF+INH+STR Resistant | 14 (28%) |
|  |  | RIF +INH+EMB Resistant | 9 (18%) |
|  |  | STR+RIF+INH+EMB Resistant | 6 (12%) |
|  |  | RIF Mono-Resistant | 1 (2%) |

Notation- Rifampicin: RIF; Isoniazid: INH; Streptomycin: STR; Ethambutol: EMB

### Minimum inhibitory concentration (MIC)

To determine the minimum inhibitory concentration (MIC), a colorimetric redox indicator (CRI) assay was performed against Rifampicin on all the 151 MTB isolates. The isolates which showed MIC value either equal to or higher than 0.50μg/mL were considered as resistant or uninhibited. For test interpretation, the color was compared to the color present in the growth control well. For quality control, the plates were also read by another laboratory member and the value which was in agreement was considered MIC for the isolate.

Out of the 151 isolates tested, 63(41.72%) had MIC of ≤0.0625µg/mL, and 18(11.92% had MIC of 0.125µg/mL. These were considered sensitive to RIF by CRI and MGIT-960 system (at critical concentration 1.0µg/mL). While 4(2.64%) isolates had MIC of 0.25µg/mL, 12(7.94%) had MIC of 0.5µg/mL, and 08(5.29%) had MIC of 1.0µg/mL These were identified as low-level RIF resistant (MIC ranging between ≥0.25µg/mL to <2.0µg/mL). Of the forty-six (30.46%) isolates, (9(5.96%) had MIC of 2µg/mL and 37(24.50%) had MIC of >2µg/mL) and were considered as high-level rifampicin resistant isolates (Figure 1).

**Figure 1.**
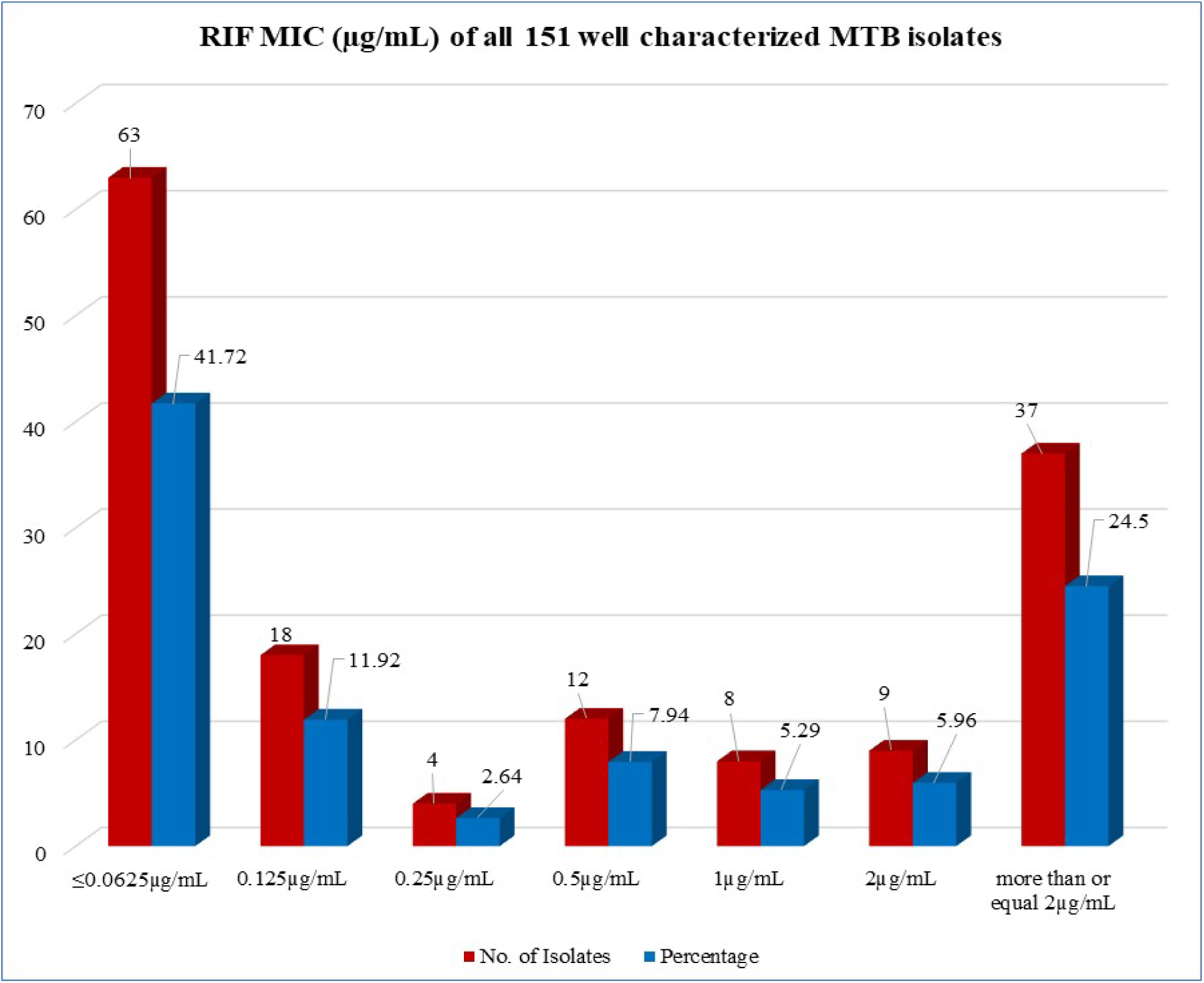
Results of the rifampicin MIC (in µg/mL) for all 151 well characterized MTB isolates.

Out of 151 isolates, 20 (13.24%) isolates had discordance for RIF susceptibility, and 04(2.6%) isolates were in concordance by MGIT960 (Table 2). All these 24 isolates were identified as having low-level RIF resistance (MIC ranging from ≥0.25µg/mL to <2.0µg/mL) by CRI assay.

**Table 2.** Results of Phenotypic RIF susceptibility tested by MGIT and CRI.

| S. No. | Isolate's RIF status using MGIT 960 | Growth status using MGIT 960 (RIF 1.0$\mu\text{g/mL}$) | RIF MIC ($\mu\text{g/mL}$) | CRI Result growth observations |
| --- | --- | --- | --- | --- |
| 1 | Resistant (4) | Delayed growth | 1.0 | Growth Present |
| 2 |  |  |  |  |
| 3 |  |  |  |  |
| 4 |  |  |  |  |
| 5 | Sensitive (20) | Growth Absent | 1.0 | Growth Present |
| 6 |  |  |  |  |
| 7 |  |  |  |  |
| 8 |  |  |  |  |
| 9 |  | Growth Absent | 0.50 | Growth Present |
| 10 |  |  |  |  |
| 11 |  |  |  |  |
| 12 |  |  |  |  |
| 13 |  |  |  |  |
| 14 |  |  |  |  |
| 15 |  |  |  |  |
| 16 |  |  |  |  |
| 17 |  |  |  |  |
| 18 |  |  |  |  |
| 19 |  |  |  |  |
| 20 |  |  |  |  |
| 21 |  | Growth Absent | 0.25 | Growth Present |
| 22 |  |  |  |  |
| 23 |  |  |  |  |
| 24 |  |  |  |  |

**Table 3:** Distribution of mutations and MIC of the isolates (n=24)

| S. No. | Mutations |  | RIF MIC (µg/mL) | No. of Isolates |
| --- | --- | --- | --- | --- |
|  | Nucleotide change | Amino acid change |  |  |
| 1 | ND | ND(4) | 0.25 | 4 |
| 2 |  |  |  |  |
| 3 |  |  |  |  |
| 4 |  |  |  |  |
| 5 | ATC-TTC | lle572Phe(5) | 0.5 | 12 |
| 6 |  |  |  |  |
| 7 |  |  |  |  |
| 8 |  |  |  |  |
| 9 |  |  |  |  |
| 10 | CAC-CTC | His526Leu(2) |  |  |
| 11 |  |  |  |  |
| 12 | GAC-TAC | Asp516Tyr(1) |  |  |
| 13 | CTG-CCG | Leu533Pro(3) |  |  |
| 14 |  |  |  |  |
| 15 |  |  |  |  |
| 16 | ND | ND(1) |  |  |
| 17 | TCG-TTG | Ser531Leu(2) | 1 | 8 |
| 18 |  |  |  |  |
| 19 | TCG-TGG, ATC-TTC | Ser531Trp, Ile572Phe(1) |  |  |
| 20 | GAC-GTC | Asp516val(1) |  |  |
| 21 | ATC-TTC | lle572Phe(2) |  |  |
| 22 |  |  |  |  |
| 23 | CAC-AAC, ATC-TTC | His526Asn, Ile572Phe(1) |  |  |
| 24 | CTG-CCG, CCT-CGT | Leu533Pro, Pro564Arg(1) |  |  |
| Total |  |  |  | 24 |
| ND: not detected |  |  |  |  |

### Sequencing Results

Of the 24 isolates, mutations in *rpoB* gene including both within RRDR and outside RRDR region were detected in 19 isolates (**Figure 2**). With 3 of these isolates having double mutation including both inside RRDR and outside the RRDR [TCG-TGG (*Ser531Trp*) + ATC-TTC (*Ile572Phe*)1; CAC-AAC (*His526Asn*) +ATC-TTC (*Ile572Phe*)1; CTG-CCG (*Leu533Pro*) +CCT-CGT(*Pro564Arg*)1]. The remaining 16 isolates possess single *rpoB* mutations. Of them 9 isolates had mutation inside the RRDR i.e., TCG-TTG (*Ser531Leu*; 2) GAC-GTC (*Asp516val*; 1), CAC-CTC (*His526Leu*; 2), GAC-TAC (*Asp516Tyr*; 1), and CTG-CCG (*Leu533Pro*; 3) while 7 isolates had ATC-TTC (*Ile572Phe*;7) mutation outside of RRDR of *rpoB* gene. The 4 isolates had no mutation.

**Figure 2.**
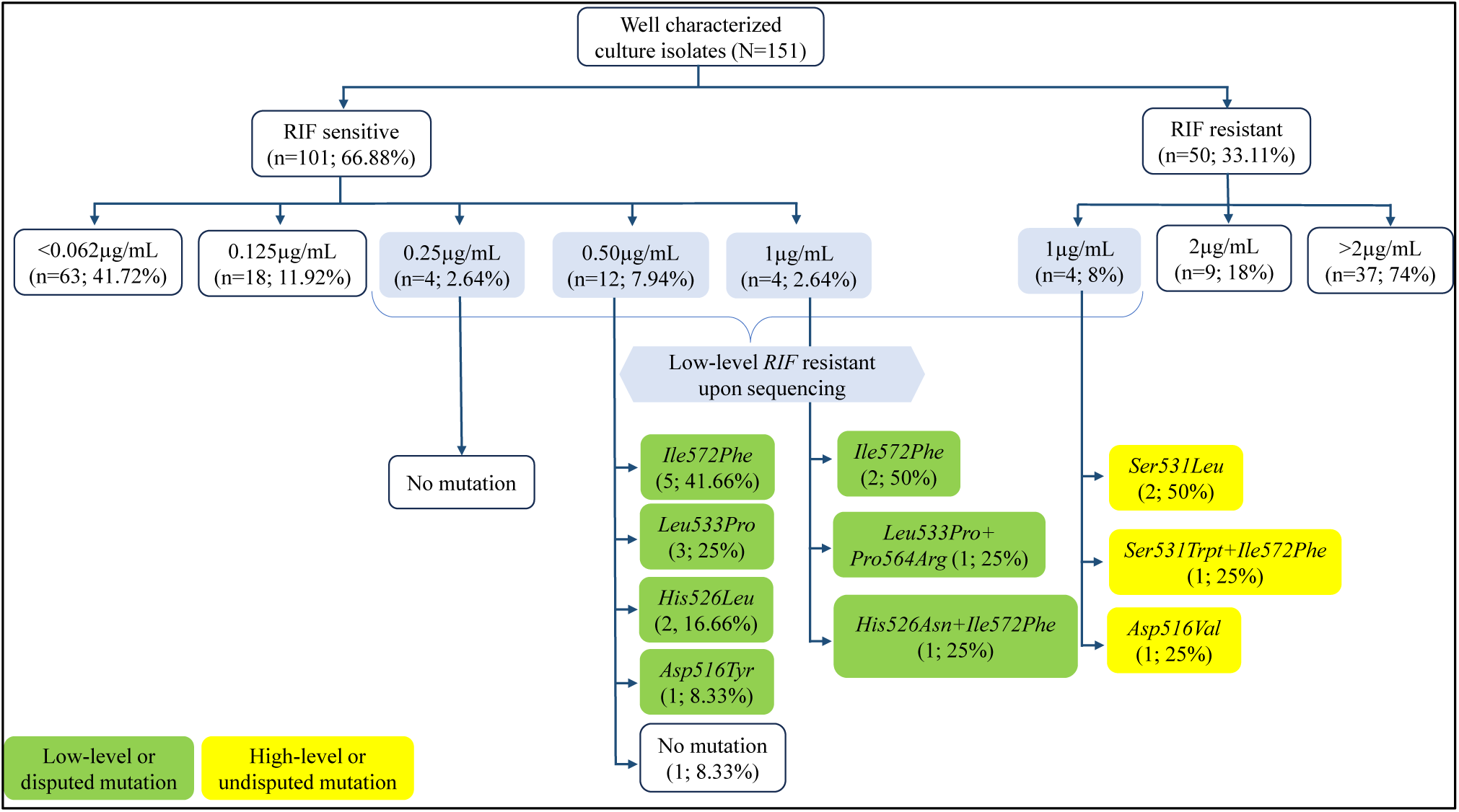
Distribution of the *rpoB* gene mutations confirmed by sequencing in the studied *M. tuberculosis* isolates (n=24)

**Figure 3.**
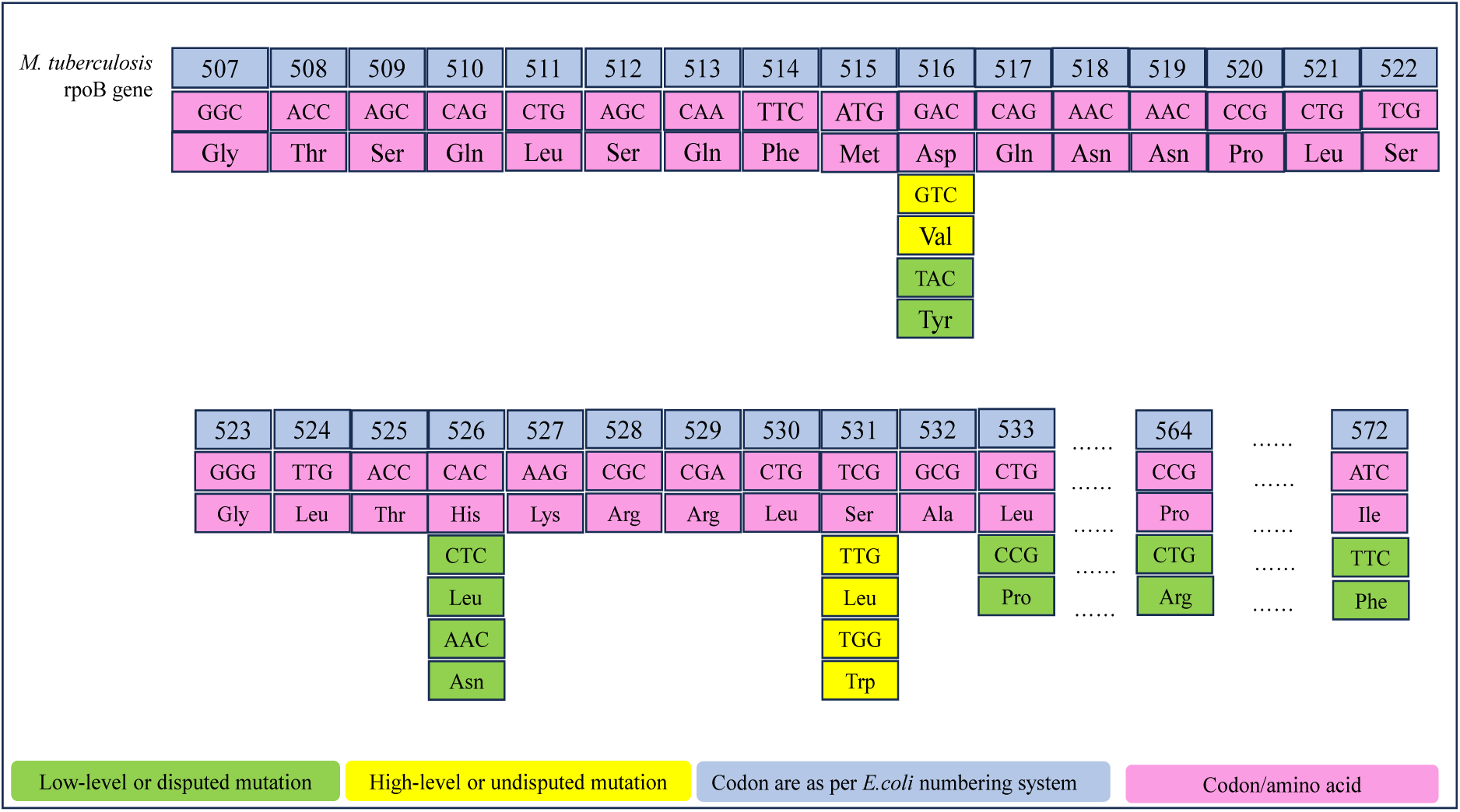
Schematic representation of codon positions of *rpoB* gene at which mutaions found in our study.

Majority of the isolates exhibiting low-level phenotypical RIF resistance were found to be associated with disputed *rpoB* mutations as follows *His526Leu,* 2; His*526Asn,*1; *Asp516Tyr*, 1; *Leu533Pro*, 4; *Ile572Phe*, 9. Although mutations associated with high-level RIF resistance *Ser531Leu* (2), *Ser531Trp* (1) and *Asp516val* (1), were also found. Sequencing results of 24 MTB isolates evaluated for RIF-resistance (CRI) are summarized in **table** (3).

## Discussion

Rifampicin resistance is considered as a surrogate marker of multi-drug resistant tuberculosis (MDR-TB). Successful treatment of MDR-TB is based on a timely and correct diagnosis and the drug susceptibility test (DST), which provides evidence for selecting an effective second line drug regimen. In the past 15 years, several molecular diagnostics methods, that allow the rapid detection of drug resistance in MTB directly from specimens, have been developed, overcoming the limitations of conventional phenotypic methods. Also, a number of these commercial molecular assays have been endorsed by WHO, which include GenoType MTBDR*plus* (Hain LifeScience, Germany), GeneXpert MTB/RIF (Cepheid, USA), and Truenat MTB (Molbio diagnostics Pvt Ltd, India). These assays detect RIF resistance conferring genetic mutations in the rifampicin resistance determining region (RRDR). However, the biggest limitation of these tests is that they miss the genetic mutations that are outside the target (RRDR) region and often considered as are disputed to confer the clinical drug resistance (11). Moreover, the molecular tests are unable to determine the level of phenotypic resistance. On the other hand, the WHO endorsed liquid culture-based phenotypic DST, which still remains the gold standard, is also not able to detect all clinically relevant resistant cases.

Using the phenotypic methods, to level an isolate as sensitive or resistance, the isolate has to be tested *in-vitro* under the drug challenge. Critical concentration (CC), which is defined as the minimum concentration of the anti-tuberculosis drug in the culture medium that is able to inhibit 99.0% growth of *M. tuberculosis*, has a crucial consideration. This concentration keeps on changing depending on the baseline resistance to a particular drug at the particular period, in that group of population. WHO recommended 1.0µg/mL as critical concentration for RIF resistance (28).

Recent studies have also shown that liquid culture systems such as MGIT-960 often fail to detect strains exhibiting low-level (eg. MIC of <1.0μg/mL) resistance to RIF (29). These low-level RIF-resistant strains with MICs below the critical concentration mostly contain specific mutations, within the hot-spot region of the rpoB gene (HSR-*rpoBI)*.

Strains with high-level resistance will have clearly elevated RIF MIC (≥16.0µg/mL) and phenotypically these are correctly identified as resistant in MGIT. Similarly strains having low-level resistance with RIF MIC between 0.25µg/mL to 1.0µg/mL will be identified phenotypically susceptible at in MGIT. Several of these isolates might have clinical resistance (29). Torrea et al, (30) reported that by using the lower level of MIC a high number of isolates could be leveled as resistant. In their study when the critical concentration was reduced to 0.50µg/mL they detected 10% additional cases and when the critical concentration was further reduced to 0.25µg/mL an additional 25.70% cases could be detected (30). Shea et. al. (2020) also reported similar results where they reported that 18.9% strains were RIF susceptible in MGIT at 1.0µg/mL, but had low-level RIF resistant by MIC testing (29). Others have also reported similar observations (11).

In our study, 15 isolates that were detected as sensitive by MGIT at standard critical concentration (1.0µg/mL) but found to exhibit low-level RIF resistance by CRI assay, were sequence-confirmed for mutations that reported to confer low-level resistance by sanger sequencing based upon the presence or absence of known *rpoB* mutations. Among them, we observed that all the isolates contained non-synonymous mutations, the commonest being the *Ile572Phe*, followed by *Leu533Pro*, *His526Leu*, *His526Asn*+*Ile572Phe*, *Asp516Tyr*, and *Leu533Pro*+*Pro564Arg* (**Figure 2**).

These are reported to be disputed mutations exhibiting low-level RIF resistance. The codon positions of *rpoB* gene at which mutations found in our study are shown in figure (**3**). However, there is no consensus that how much reduction in the presently WHO recommended critical concentration of 1.0µg/mL can be done, or should it be changed at all. The WHO Technical Expert Group in 2020 deliberated on this issue and agreed that the critical concentration of 0.50µg/mL will be idealistic, but it could not verify with CLSI, if there is sufficient evidence to recommend lowering of the critical concentration of RIF from 1.0µg/mL to 0.50µg/mL in MGIT culture-based drug susceptibility testing (3). WHO is also considerate that if the critical concentration of RIF is reduced from 1.0µg/mL to 0.50µg/mL in MGIT culture-based drug susceptibility testing, the number of RIF resistant cases, which is also the surrogate marker of MDR-TB, will increase exponentially. It may not be feasible for the national TB control programs to handle such a large number of resistant cases, which are hitherto are considered to be susceptible using the currently recommended critical concentration of 1.0µg/mL.

The current guidelines of WHO, are mostly reliant on molecular detection of resistance conferring mutations, because the molecular tests are rapid and less cumbersome and can be done directly on the clinical samples without even culturing the pathogen. However, these molecular tests including the GeneXpert MTB/RIF and LPA have limited scope of detecting the genetic mutations in a specific (hot spot) area of *rpoB* gene. Any mutation outside the hotspot (e.g., *Ile572Phe*) or on the extreme ends of RRDR region will be missed and these patients will be treated with standard regimen of drug susceptible TB cases. While actually, these patients need to be treated with an MDR-TB regimen (15,11,31).

The distribution of these mutations seen in our study are in line with previous studies (30,11), but the frequencies of various mutations were different. In our study, we found that *Ile572Phe* mutation was commonest while Torrea et al (30) found that *Asp516Tyr* was commonest while Rigouts et al (11) found that *His526Leu* was commonest mutation when using the MIC of 0.50µg/mL (30). In a recent report (32), out of 308 isolates 227 isolates were found to be RIF-susceptible by culture-based DST, but 8(3.52%) of them were identified as resistant by MTBDR*plus*. On WGS, they found that 2 isolates had *Leu533Pro* mutations, another 2 had *Asp516Tyr*, and one each had *His526Leu*, *Leu511Pro*, *His526Asn* mutations. The MICs of these isolates ranged from 0.125μg/mL to 1.0μg/mL. We did not find any of these mutations at MIC ≤0.25μg/mL.

In our study, 20% isolates were found to have *Leu533Pro* mutation at MIC of 0.5µg/mL which is highest frequency reported so far. Rando-Segura et al.(32), from Angola reported a frequency of 12.50%, Torrea et al (30) from Belgium reported 11.11%, and Salaam-Dreyer et al from South Africa reported the frequency of 8% (33) only. While Berrada et al from USA reported very low frequency of 2.3% only (34). At MIC of 0.50µg/mL, we also found that *His526Leu* mutation was present in 13.33% isolates. The frequency of this mutation was much higher in our isolates as compared to other studies, where the frequency ranged from as low as 2.4% to 12.5% (30–34).

We found frequency of mutation *Asp516Tyr*, *His526Asn*+*Ile572Phe* and *533Pro*+*Pro564Arg* very low at MIC values between 0.5µg/mL and 1µg/mL. Others have also reported low frequency of these mutations, except Andres et al, from Germany who reported prevalence of 25% (11,34,35).

At MIC of 1.0µg/mL we found that *Ile572Phe* mutation was commonest (13.33%). This frequency falls within the range reported by other workers (29,33). The *His526Leu* mutation was found in our isolates at this drug concentration, but others have reported its frequency up to 12.5% (32). We could detect *Leu533Pro* mutation in 6.6% of our isolates, while Ho et al, from Australia reported prevalence of 20% and Wang et al, from China reported frequency of this mutation in 3.3% isolates (18,36).

Our study also showed novel findings of dual mutations as mentioned above. Such dual mutations have not been reported by anyone so far.

As such not many studies have been done on this aspect to correlate the MIC and mutations but even very few on the low-level resistance conferring mutations. There is only one study from India (37). Of the 84 RIF resistant isolates, they found 16.66% isolates having disputed mutations, at MIC of 0.25 to 0.5µg/mL, *Asp516Tyr* in 1.2%, and *His526Asn* in 2.38%, respectively. Therefore, ours becomes the unique study in which we have compared the frequency of various mutations often considered to be disputed or low resistance conferring with the low MIC in the MGIT culture. Our study is not only first of its kind from India but also from entire south-east Asia to best of knowledge and literature search till the writing of this manuscript.

A number of recent studies have argued in favour of lowering the critical concentration. Yip et al (2006) from Hong-Kong proposed that strains carrying *Leu511Pro*, *Leu533Pro*, and *His526Leu* mutations represented almost 22% (19/85) of all the mutations compared to less than 10% among all phenotypically RIF-resistant strains of previous years, meaning that approximately 12% phenotypically RIF susceptible isolates (38). They also mentioned that the frequency of these mutations may vary from region to region. This further poses a question, if the frequency of these mutations also varies in various genotypes (39).

From our study and from others’ work, it is evident that lowering of the critical concentration in phenotypic methods and inclusion of the above-mentioned mutations and their probes in currently used molecular methods can detect several such cases which are hitherto considered as drug susceptible and treatment with primary anti-TB drugs is initiated.

Our data shows that a significant number of isolates carry low-level RIF resistance and the patient carrying these strains are falsely levelled as susceptible, and thereby treated with primary anti-TB drugs. Many of these patients might later on manifest as MDR cases. This will be worthwhile to conduct nationwide research to find these mutations and follow these patients who carry these so-called low-level or disputed mutations, to see what percentage of these patients actually get cured using the primary antitubercular disease (ATD). This evidence is necessary before advising lowering of the critical concentration and including these mutations in the currently used molecular tests, because this might pose a huge human resource and economic burden on the national TB elimination/control programs.

We are aware that impromptu lowering of the critical concentration in DST, might pose a huge human resource and economic burden on the national TB elimination/control programmes. Hence, we propose that it will be worthwhile to conduct nationwide research to detect these mutations in the RIF susceptible patients and follow these patients to see how many of these patients get cured with primary anti-TB drugs. We hypothesis that a significant number of patients carrying low-level drug resistance mutation will become clinically drug resistant.

## Conclusion

Comparing the MIC and genetic mutations within the HSR-*rpoB* gene region of *M. tuberculosis*, with the results of standard WHO endorsed MGIT 960 drug susceptibility testing system which uses critical concentration of 1.0µg/mL for rifampicin, we observed that a significant number of clinical isolates shown as sensitive by this system carry a low-level rifampicin resistance and specific genetic mutations. This means that a significant number of patients are put on an anti-tuberculosis drug regimen, which is not effective, and therefore, the crucial concentration needs to be reviewed.

We are aware that impromptu lowering of the critical concentration in DST, might pose a huge human resource and economic burden on the national TB elimination/control programmes. Hence, we propose that it will be worthwhile to conduct research to detect low level drug resistance conferring mutations using novel diagnostic methods and follow these patients to see what percentage of these patients actually get cured with primary anti-TB drugs and how many require second line regimen. We hypothesis that a significant number of patients carrying low-level drug resistance mutation will become clinically drug resistant.

### Funding

This study was supported by project grant No.(5/8/5/41/2016/ECD-I) funded by the Indian Council of Medical Research, New Delhi.

### Conflict of Interest

None declared

